# Chemical Coverage of the Human Reactome

**DOI:** 10.1101/2024.04.23.590739

**Authors:** Haejin Angela Kwak, Lihua Liu, Claudia Tredup, Sandra Röhm, Panagiotis Prinos, Jark Böttcher, Matthieu Schapira

## Abstract

Chemical probes and chemogenomic compounds are valuable tools to link gene to phenotype, explore human biology and uncover novel targets for precision medicine. A growing federation of scientists is contributing to the mission of Target 2035 – discovering chemical tools for all druggable human proteins by the year 2035. It is expected that these compounds will enable the understanding of the regulation of cellular machineries and biological processes across the compendium of signaling pathways that animate cellular life. Here, we draw a landscape of the current chemical coverage of the human Reactome. We find that even though available chemical probes and chemogenomic compounds are targeting only 3% of the human proteome, they cover 53% of the human Reactome, due to the fact that 46% of human proteins are involved in more than one cellular pathway. As such, existing chemical probes and chemogenomic compounds already represent a versatile toolkit to manipulate a vast portion of human biology. Pathways targeted by existing drugs may be enriched in unknown but valid drug targets and could be prioritized in future Target 2035 efforts.

## Introduction

Twenty three years after the sequencing of the human genome, our knowledge of the human proteome remains incomplete.^1–3^ CRISPR-based genome targeting and editing enables the functional characterization of proteins and is broadly applied in target prioritisation for pharmaceutical intervention.^4,5^ A complementary approach, differing both in mode of interference and applicability, is to target a protein with chemical tools or antibodies.^6,7^ A key obstacle in studying the function of underexplored proteins is the lack of high-quality chemical tools.^1–3^ To accelerate the characterization of the human proteome, biomedical scientists around the globe have formed a federation called Target 2035 to develop and openly provide rigorously characterized chemical tools, such as chemical probes and chemogenomic compounds, for the entire druggable human proteome by year 2035.^2,3,8^ Target 2035 contributors include, but are not limited to, the Structural Genomics Consortium (SGC)^9^, EUbOPEN^10^, RESOLUTE^11^, NCATS at NIH^12^, Boehringer Ingelheim’s opnMe^13^, and portals such as Chemical Probes Portal^14^, Pharos^15^, Probes and Drugs^16^, and ChemBioPort^17^.

Chemical probes are selective and potent small-molecule modulators of proteins that can be valuable in biochemical, cell-based, or animal models.^18^ They are used to link a phenotype to a target protein and ideally bind to and engage a single target. Chemogenomic compounds are highly annotated molecules within libraries of congeneric compounds that are used to investigate members of a certain protein family.^19,20^ They are less selective than chemical probes, but the effects of multiple chemogenomic compounds with overlapping selectivity profiles can be compared and used to decouple their polypharmacology and link a protein to a phenotype.^21^

Here we analyze Target 2035’s current chemical coverage of proteins, protein families, and biological pathways. Evaluating the progression by protein or by protein family reveals how much of the human proteome, kinome and GPCRome for instance, is targeted by chemical tools. Monitoring the progression by pathway highlights areas of human biology that are underexplored by the chemical biology community and provides a new metric to prioritize the development of chemical tools.

## Results

### Chemical Coverage of the Human Proteome

To track Target 2035’s progression by protein, we analyzed the coverage of the human proteome by chemical probes, chemogenomic compounds, and drugs (Figure 1a). Out of the roughly 20,000 human proteins that constitute the human proteome^39^, we find that 2,331 (∼11.7%) are targeted by a chemical tool and/or drug. This includes 437 (∼2.2%) targeted by chemical probes, 353 (∼1.8%) by chemogenomic compounds, and 2,112 (∼10.6%) by drugs (Table S1). We observed a significant overlap between the different compound types; for instance, only 219 of the 627 proteins targeted by chemical tools are not targeted by drugs.

**Figure 1.**
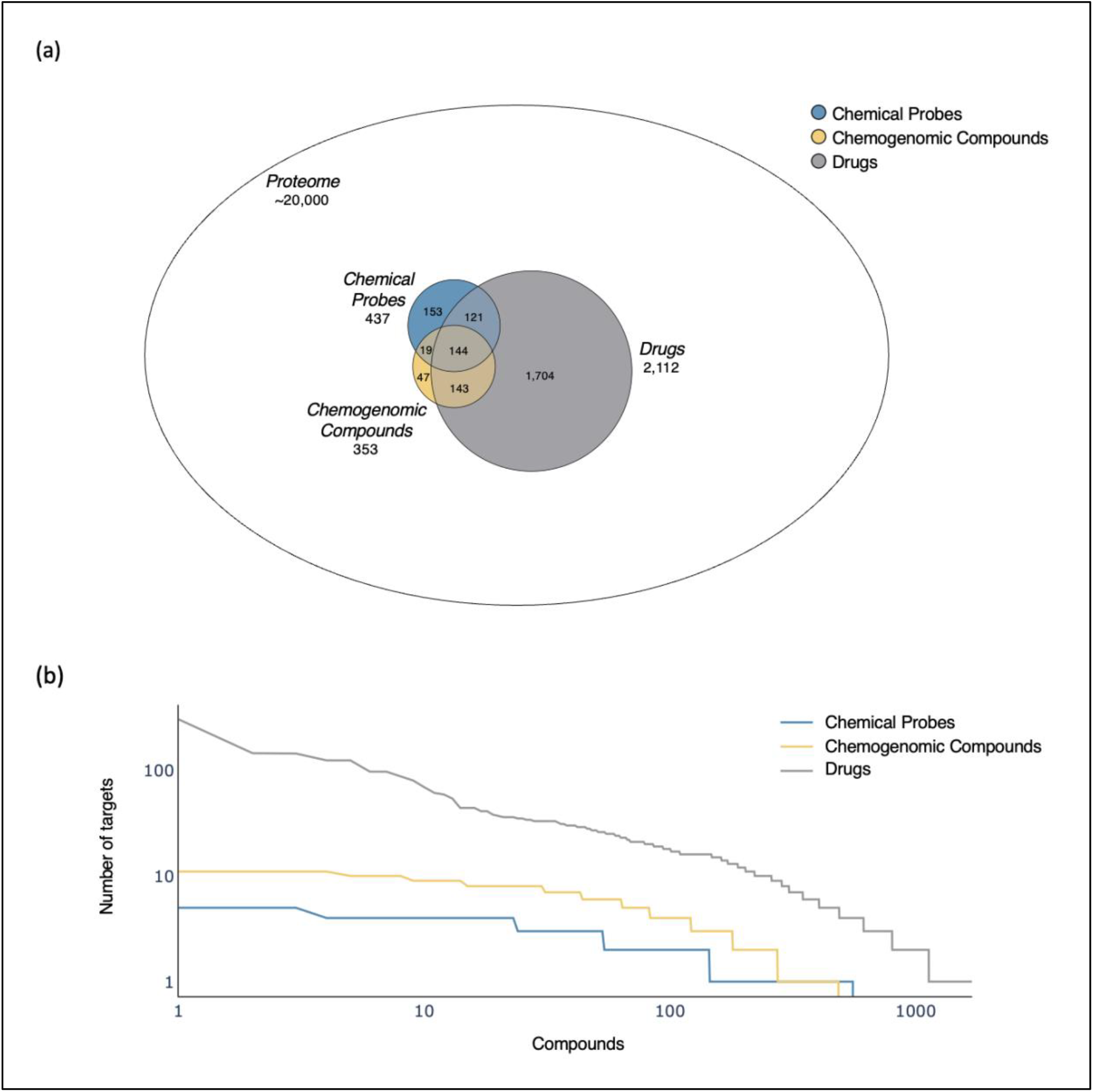
(a) Coverage of the human proteome by chemical probes, chemogenomic compounds, and drugs. Annotated are the total number of protein targets for each class of compounds. (b) Number of known targets for each of the 554 chemical probes, 484 chemogenomic compounds, and 1693 drugs.

The higher coverage by drugs can be attributed to i) their larger number of compounds (1693) compared to 554 chemical probes and 484 chemogenomic compounds, and ii) their lower selectivity (Figure 1b). Indeed, unlike chemical probes and chemogenomic compounds, high selectivity is not mandatory for drugs, many of which exert their clinical effects through polypharmacology.^40^ The number of known targets for drugs ranges from 1-305 while the number of targets for chemical probes and chemogenomic compounds fall within a much narrower range of 1-5 and 1-11 respectively. Furthermore, 74.0% of chemical probes and 43.6% of chemogenomic compounds have a single target, compared to 33.4% of drugs, which is likely an underestimate as the proteome-wide selectivity profile of drugs is generally unknown.^18^

The difference in selectivity between the different compound types reflects their distinct purposes. Chemical probes and chemogenomic compounds are tool compounds used to answer specific biological questions. On the other hand, the purpose of drugs is to achieve therapeutic benefit rather than to study a protein, therefore, if the benefit outweighs the risk, the lack of selectivity of a drug is acceptable. For example, fostamatinib is a promiscuous drug with 305 known targets that has been approved by the FDA as the first-in-class spleen tyrosine kinase inhibitor for the treatment of chronic immune thrombocytopenia.^41,42^

### Chemical Coverage of Reactome Pathways

To track Target 2035’s progression by biological pathways, we mapped chemical probes and chemogenomic compounds onto human pathways from the Reactome knowledgebase.^37^ Specifically, we determined the total number of proteins in each pathway and the number of those proteins targeted by chemical probes and chemogenomic compounds (Figure 2 and Table S2). A similar analysis was conducted with drugs as a reference. The human Reactome knowledgebase contains 2673 pathways, with 29 of them being top-level pathways, under which the remaining 2644 are hierarchically organized in a parent-child manner.^38^ When a protein is present in a child pathway, it is also present in its parent pathway in a recursive manner until a top-level pathway is reached.^43^

**Figure 2.**
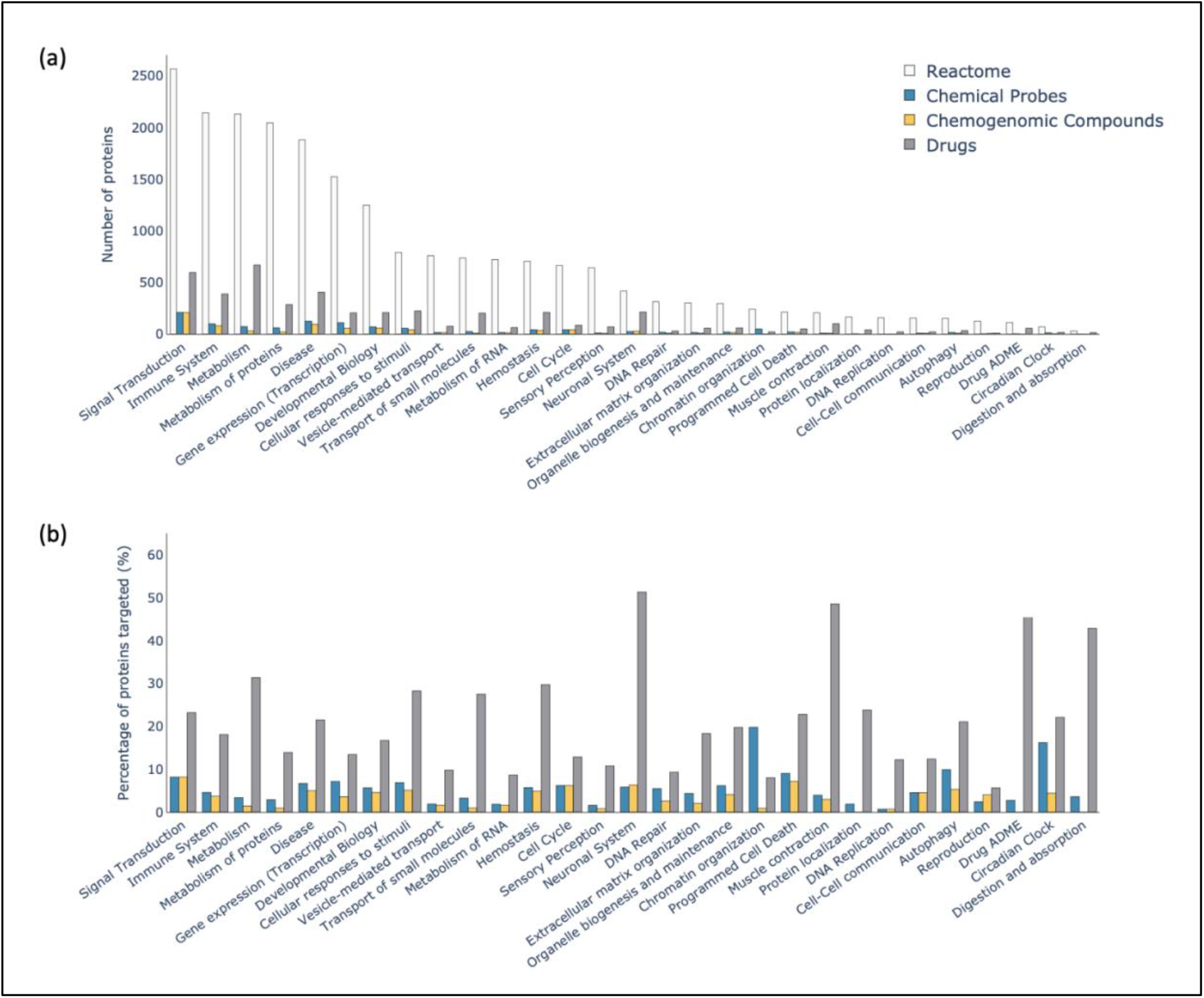
Chemical coverage of the top-level pathways of Reactome. (a) Total number of proteins and number targeted by chemical probes, chemogenomic compounds, and drugs for each top-level pathway. (b) Percentage of proteins targeted.

Strikingly, only five top-level pathways have a coverage by chemical tools that exceeds 10%, compared to 24 by drugs (Figure 2b, Table S2). Moreover, multiple pathways have a relatively high percentage of proteins targeted by drugs, including *neuronal system* (51.3%), *muscle contraction* (48.5%) and *metabolism* (31.3%). We also note that drugs cover a relatively lower percentage of the *disease* pathway (21.5%), but a closer inspection reveals great heterogeneity amongst its child pathways; for example, low number of proteins and high coverage in *disease of the neuronal system* where there are drugs for 6 out of 13 proteins, which contrasts with high number of proteins and low coverage in *infectious disease* where there are drugs for 211 out of 1078 proteins (Figure S1a and Table S2). The only top-level pathway that is better covered by chemical probes than drugs, 19.8% versus 8.0% respectively, *is chromatin organization* (Figure 2b and Table S2). The good coverage is driven, among others, by chemical probes targeting readers of lysine methylation^44^, acetylation^45^, and protein methyltransferases^46^. The top-level pathway best covered by chemogenomic compounds is *signal transduction* (8.1%) (Figure 2b and Table S2), as expected bearing in mind the current efforts to develop chemogenomic compound sets for the protein kinase and GPCR families.^2,3,47–49^

The lowest level (“most granular”) pathways of Reactome, ones not subdivided into child pathways, depict the true hubs of the proteome where the molecular interactions and reactions take place. Out of the 1883 lowest level pathways, 869 (46.1%) are targeted by a chemical probe and 689 (36.6%) are targeted by a chemogenomic compound (Figure 3a and Table S2). This is in striking contrast with the ∼3.1% coverage of the human proteome by chemical probes and chemogenomic compounds combined (Figure 1a). The main reason for this apparent disconnect is that one protein can be involved in multiple pathways. Indeed, 5,128 out of the 11,175 (45.9%) proteins in Reactome are found in more than one lowest level pathway, including ribosomal protein S27a, which spans as much as 165 (Figure 3b). Even at the top-level pathways, the specificity of these proteins remains poor with 4,605 out of the 11,175 (41.2%) proteins in Reactome being found in more than one top-level pathway (Figure 3d). These results indicate that even highly selective chemical tools may not achieve ideal pathway specificity. Conversely, the compendium of available chemical probes and chemogenomic compounds allows for the pharmacological targeting of 1,556 out of 2,673 pathways, or 52.8% of the human Reactome (Table S2), and already constitutes a versatile toolkit to investigate cellular machineries across a vast portion of human biology. As the chemical coverage of the proteome increases, it may become possible to compare the cellular effect of multiple chemical tools with distinct but overlapping pathway specificity profiles to link pathway to phenotype. Conversely, the association of a phenotype to a pathway will be stronger if multiple chemical tools that target different nodes within a pathway all induce the same phenotype.

**Figure 3.**
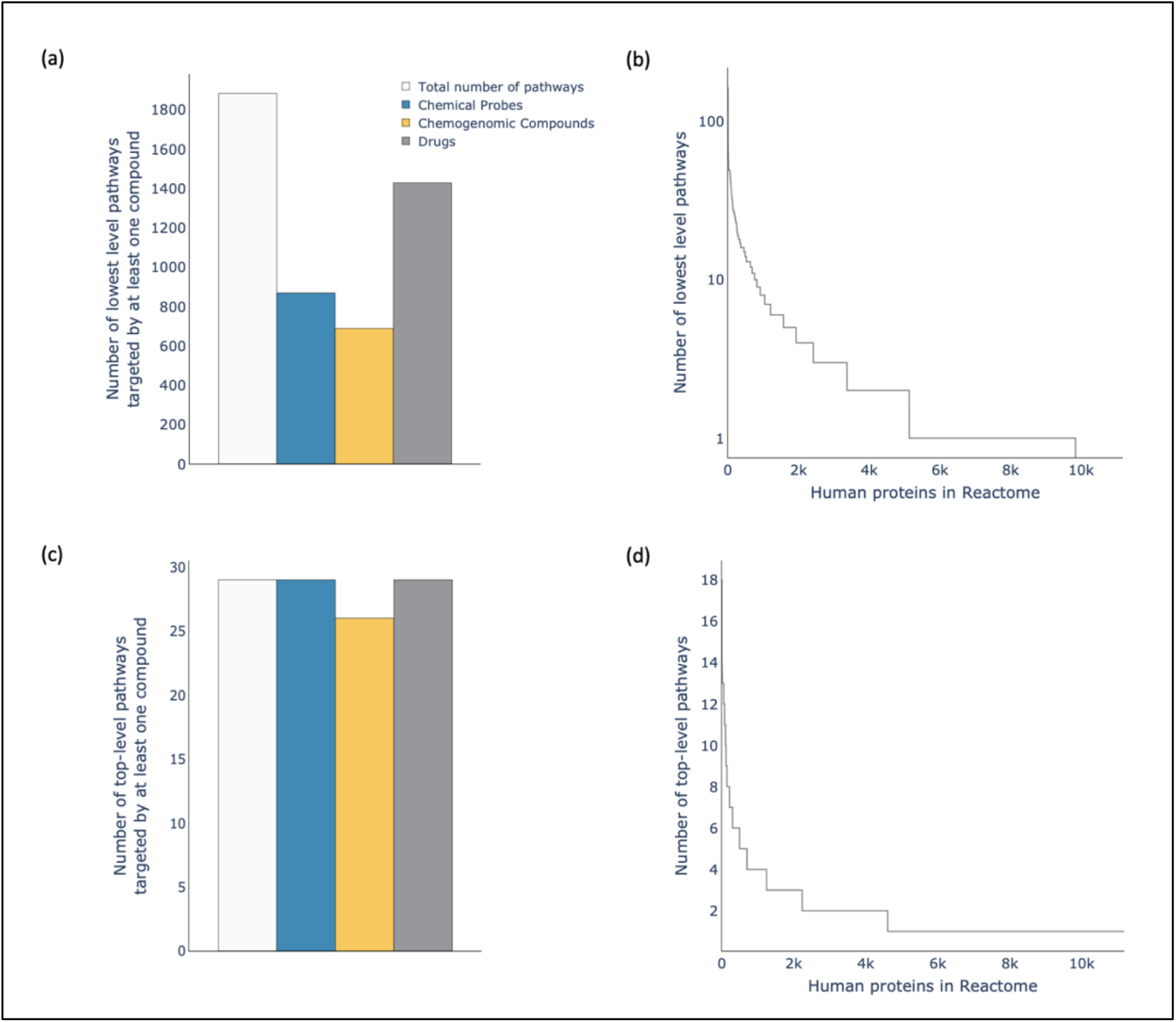
The number of (a) lowest level pathways and (c) top-level pathways of Reactome that are targeted by at least one compound for each chemical probes, chemogenomic compounds, and drugs. The number of (b) lowest-level pathways and (d) top-level pathways that each human protein of Reactome resides in.

### Chemical Coverage of the Kinome and GPCRome

We next focused our analysis on kinases and GPCRs – two highly populated and well established classes of therapeutic targets with high chemical coverage and dedicated chemical tool development efforts from the Target 2035 community.^2,3,47–49^ As expected, we find high chemical coverage for these protein families, with 67.7% of the 538 human kinases and 20.3% of the 802 human GPCRs having at least one reported small molecule compound, compared with ∼11.7% of the human proteome. Kinases are significantly more targeted by chemogenomic compounds (49.8%) and drugs (58.0%) than by chemical probes (27.7%) (Figure 4a and Table S3), but it should be noted that the especially high coverage by drugs mostly reflects the poor selectivity of some of these molecules. For example, the drug fostamatinib targets 274 kinases.^31^ In spite of the relatively high chemical coverage of kinases, we note that 243 (45.2%) are targeted neither by chemical probes, chemogenomic compounds, nor a selective drug (which we loosely define as a drug with less than 10 known targets) (Table S3), which supports current efforts to chemically explore the dark kinome.^47,48,50^ We also find that, like the kinome, the GPCRome is more targeted by drugs (17.3%) than chemical probes (7.0%) or chemogenomic compounds (7.6%) (Figure 4a and Table S4), but annotations from DrugBank indicate that GPCR ligands are more selective than kinase inhibitors. Indeed, the most promiscuous molecule, aripiprazole, targets 34 GPCRs, compared with fostamatinib that targets 274 kinases as mentioned above.^31^

**Figure 4.**
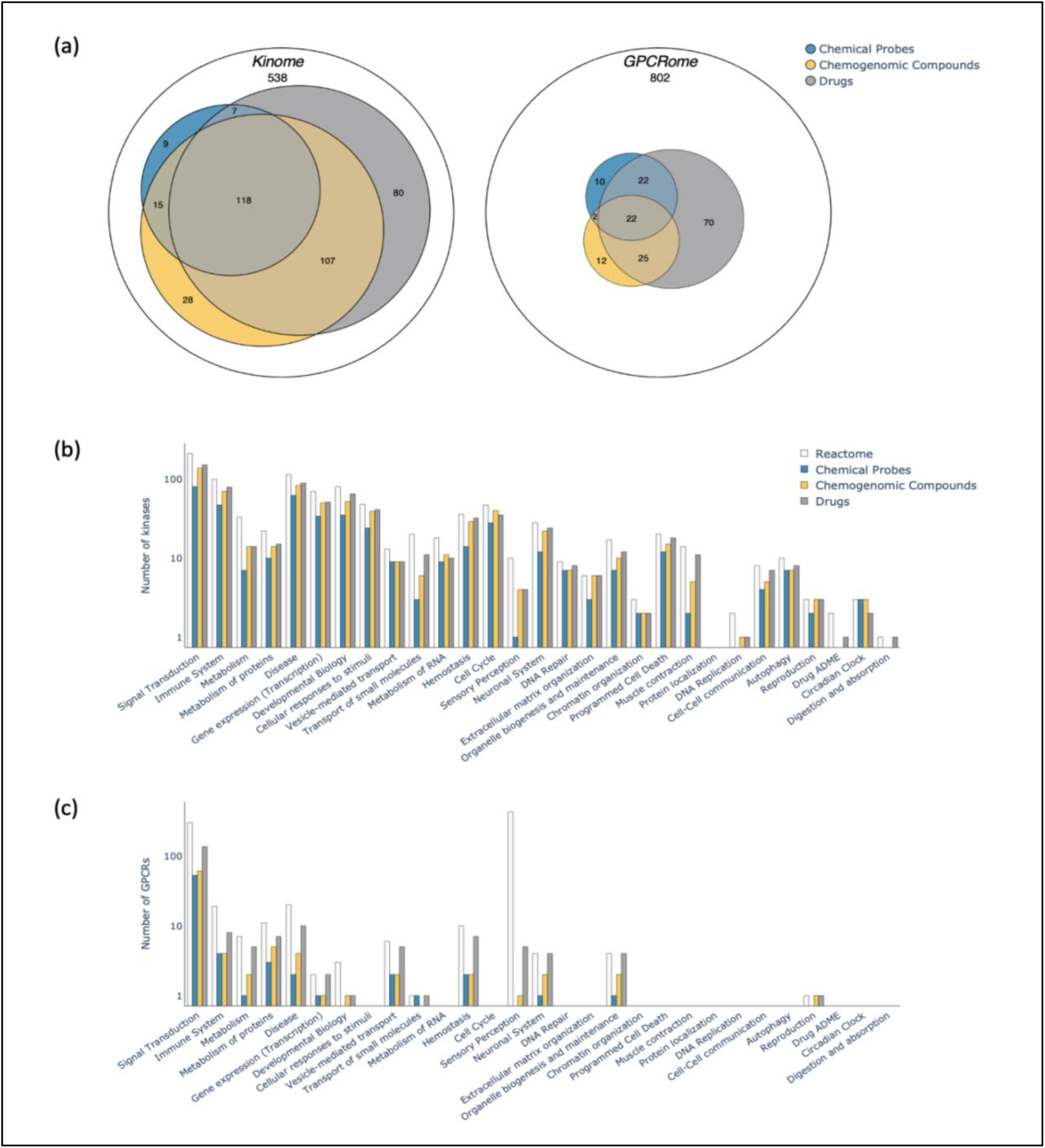
Chemical coverage of kinases and GPCRs. (a) Coverage of the human kinome and GPCRome by chemical probes, chemogenomic compounds, and drugs. (b,c) For each pathway, the total number of proteins and number targeted by chemical probes, chemogenomic compounds and drugs is shown for kinases (b) and GPCRs (c).

We next investigated the distribution of human kinases and GPCRs and their ligands across the Reactome. Kinases are found in 28 out of the 29 top-level pathways, whereas GPCRs are found in 14 (Figure 4b and 4c), reflecting that kinases are more broadly involved in biological pathways compared to GPCRs. Most kinases are found in the *signal transduction* top-level pathway as expected, and the coverage of kinases across the Reactome is generally high (Figure 4b and Table S5). A great proportion of GPCRs are found in the *sensory perception* top-level pathway reflecting the fact that more than half of the GPCRs (407) encoded by the human genome are olfactory receptors (Table S4).^36^ However, only 5 GPCRs found in this pathway are targeted by chemical tools and/or drugs, probably because sensory perception is rarely of interest in drug discovery. 301 of the remaining 395 non-olfactory GPCRs act as signal transducers in the *signal transduction* pathway (Figure 4c and Table S6). Aberrant signal transduction is of significance in clinical research and drug discovery, which is reflected by the relatively higher coverage of GPCRs by drugs (45.2%) in the *signal transduction* top-level pathway. Here again, chemical probes and chemogenomic compounds follow a similar trend of relatively higher coverage; 17.6% and 20.3% respectively (Figure 4c and Table S6). Considering the general disease association of GPCRs involved in signal transduction and the value of chemical probes and chemogenomic compounds for the exploration of unknown biology and discovery of new therapeutic targets, a strong focus on developing chemical tools for some of these poorly characterized targets in *signal transduction* is warranted. Yet, we note that only 27 of the 301 *signal transduction* GPCRs that are not targeted by a selective drug are targeted by chemical probes or chemogenomic compounds. This strongly emphasizes the need for continued effort by the Target 2035 community to focus on this protein family.

## Concluding remarks

Analyzing the coverage of the human proteome by chemical probes and chemogenomic compounds reveals that there is still a long way to go before all human proteins are targeted with a chemical tool. *With so many proteins left to target, where should the Target 2035 community prioritize their effort?* A primary focus is on the dark proteome to reveal novel molecular machineries and unexpected disease associations. Indeed, the chemical tools included in our analysis target 88 poorly characterized proteins that are yet to be annotated in the human Reactome (Table S2). A second strategy of Target 2035 is to focus on large protein families. This is in principle operationally more efficient since protein production and purification, assay development, and structural chemistry for these target classes are relatively well established, allowing for easier chemical tool development. The two strategies mentioned can be combined; for example, bringing a focus on the dark kinome or the dark GPCRome, which is particularly relevant considering the high disease association of these target classes.^51,52^

Here, we propose a third strategy – a complementary and pathway-based approach to guide Target 2035 efforts. In most cases, enough is known about human proteins to map them onto cellular pathways, but their disease association and potential as a therapeutic target remain unclear. When a protein is mapped onto pathways enriched with drug targets, their likelihood of being a (so far unknown) drug target is increased. For instance, 118 out of 302 GPCRs playing a role in *signal transduction* are targeted by selective drugs (Table S6), clearly highlighting this subfamily to be enriched with therapeutic targets. Chemical probes and chemogenomic compounds would be valuable tools to reveal which of the remaining 184 are also potential drug targets, but they are only available for 27. This highlights a need for more chemical tools targeting GPCRs involved in *signal transduction*. Similarly, other areas of the Reactome, such as *disease* or *immune system*, are enriched in drugs but lacking in chemical tools (Figure 2) and appear to be good candidates where future chemical tools could reveal novel therapeutic targets. However, it is also important not to overlook the development of chemical tools for underexplored pathways, ones with low chemical coverage, to expand our understanding of human biology.

While the chemical coverage of the human proteome remains far from complete, achieving this ambitious goal by year 2035 remains possible thanks to the continuous scientific, technical, and logistical progress of the Target 2035 community.^2,3,8^ To accelerate the clinical impact of this global effort, we propose that a focus should be brought on proteins involved in biological pathways enriched in drugs and deprived of chemical tools, as was highlighted in this analysis. Another valid but longer-term strategy is to focus on chemically underexplored pathways, ones so far devoid of both drugs and chemical tools. To facilitate these strategies, we provide Probe my Pathway (PmP) – a webtool available at apps.thesgc.org/pmp/ where users can navigate the coverage of the human Reactome by chemical tools.

## Acknowledgments

M.S. gratefully acknowledges support from the Canadian Strategic Innovation Fund (SIF Stream 5), NSERC (RGPIN-2019-04416), and CIHR (202309PJT). H.A.K. is supported by the CIHR Canada Graduate Scholarship – Master’s Awards. The Structural Genomics Consortium is a registered charity (no: 1097737) that receives funds from Bayer AG, Boehringer Ingelheim, Bristol Myers Squibb, Genentech, Genome Canada through Ontario Genomics Institute [OGI-196], EU/EFPIA/OICR/McGill/KTH/Diamond Innovative Medicines Initiative 2 Joint Undertaking [EUbOPEN grant 875510], Janssen, Merck KGaA (aka EMD in Canada and US), Pfizer and Takeda.

## References

1. Edwards AM, Isserlin R, Bader GD, Frye SV, Willson TM, Yu FH. Too many roads not taken. Nature. 2011;470(7333):163–165. doi:10.1038/470163a

2. Carter AJ, Kraemer O, Zwick M, Mueller-Fahrnow A, Arrowsmith CH, Edwards AM. Target 2035: probing the human proteome. Drug Discov Today. 2019;24(11):2111–2115. doi:10.1016/j.drudis.2019.06.020

3. Müller S, Ackloo S, Al Chawaf A, et al. Target 2035 - update on the quest for a probe for every protein. RSC Med Chem. 2022;13(1):13–21. doi:10.1039/d1md00228g

4. Behan FM, Iorio F, Picco G, et al. Prioritization of cancer therapeutic targets using CRISPR-Cas9 screens. Nature. 2019;568(7753):511–516. doi:10.1038/s41586-019-1103-9

5. Shi J, Wang E, Milazzo JP, Wang Z, Kinney JB, Vakoc CR. Discovery of cancer drug targets by CRISPR-Cas9 screening of protein domains. Nat Biotechnol. 2015;33(6):661–667. doi:10.1038/nbt.3235

6. Frye SV. The art of the chemical probe. Nat Chem Biol. 2010;6(3):159–161. doi:10.1038/nchembio.296

7. Workman P, Collins I. Probing the probes: fitness factors for small molecule tools. Chem Biol. 2010;17(6):561–577. doi:10.1016/j.chembiol.2010.05.013

8. Ackloo S, Antolin AA, Bartolome JM, et al. Target 2035 – an update on private sector contributions. RSC Med Chem. 2023;14(6):1002–1011. doi:10.1039/D2MD00441K

9. SGC. SGC. Accessed March 21, 2024. https://www.thesgc.org/

10. Homepage | EUbOPEN. Accessed March 21, 2024. https://www.eubopen.org/

11. RESOLUTE. RESOLUTE. Accessed March 21, 2024. https://re-solute.eu/

12. National Center for Advancing Translational Sciences | National Center for Advancing Translational Sciences. Accessed March 21, 2024. https://ncats.nih.gov/

13. Gollner A, Köster M, Nicklin P, et al. opnMe.com: a digital initiative for sharing tools with the biomedical research community. Nat Rev Drug Discov. 2022;21(7):475–476. doi:10.1038/d41573-022-00071-9

14. Antolin AA, Sanfelice D, Crisp A, et al. The Chemical Probes Portal: an expert review-based public resource to empower chemical probe assessment, selection and use. Nucleic Acids Res. 2023;51(D1):D1492–D1502. doi:10.1093/nar/gkac909

15. Kelleher KJ, Sheils TK, Mathias SL, et al. Pharos 2023: an integrated resource for the understudied human proteome. Nucleic Acids Res. 2023;51(D1):D1405–D1416. doi:10.1093/nar/gkac1033

16. Skuta C, Popr M, Muller T, et al. Probes & Drugs portal: an interactive, open data resource for chemical biology. Nat Methods. 2017;14(8):759–760. doi:10.1038/nmeth.4365

17. Liu L, Rovers E, Schapira M. ChemBioPort: an online portal to navigate the structure, function and chemical inhibition of the human proteome. Database J Biol Databases Curation. 2022;2022:baac088. doi:10.1093/database/baac088

18. Arrowsmith CH, Audia JE, Austin C, et al. The promise and peril of chemical probes. Nat Chem Biol. 2015;11(8):536–541. doi:10.1038/nchembio.1867

19. Chemogenomics. Published online January 1, 2007:921–937. doi:10.1016/B0-08-045044-X/00113-9

20. Jones LH, Bunnage ME. Applications of chemogenomic library screening in drug discovery. Nat Rev Drug Discov. 2017;16(4):285–296. doi:10.1038/nrd.2016.244

21. Dafniet B, Cerisier N, Boezio B, et al. Development of a chemogenomics library for phenotypic screening. J Cheminformatics. 2021;13(1):91. doi:10.1186/s13321-021-00569-1

22. UK C org. Welcome to the Chemical Probes Portal. Chemical Probes Portal. Accessed February 6, 2024. https://www.chemicalprobes.org/

23. https://www.opnme.com. opnMe | Boehringer Ingelheim Open Innovation Portal. https://opnme.com. Accessed December 12, 2023. https://www.opnme.com/node/1226

24. Tredup C, Ndreshkjana B, Schneider NS, et al. Deep Annotation of Donated Chemical Probes (DCP) in Organotypic Human Liver Cultures and Patient-Derived Organoids from Tumor and Normal Colorectum. ACS Chem Biol. 2023;18(4):822–836. doi:10.1021/acschembio.2c00877

25. Donated chemical probes | SGC. Accessed February 6, 2024. https://www.thesgc.org/donated-chemical-probes

26. Wells CI, Al-Ali H, Andrews DM, et al. The Kinase Chemogenomic Set (KCGS): An Open Science Resource for Kinase Vulnerability Identification. Int J Mol Sci. 2021;22(2):566. doi:10.3390/ijms22020566

27. KCGSv2.0 Data. SGC-UNC. Accessed December 13, 2023. https://www.sgc-unc.org/kcgs-data

28. Chemogenomic set | EUbOPEN. Accessed December 13, 2023. https://www.eubopen.org/chemogenomics/chemogenomic-set

29. Wishart DS, Knox C, Guo AC, et al. DrugBank: a knowledgebase for drugs, drug actions and drug targets. Nucleic Acids Res. 2008;36(Database issue):D901–906. doi:10.1093/nar/gkm958

30. Wishart DS, Feunang YD, Guo AC, et al. DrugBank 5.0: a major update to the DrugBank database for 2018. Nucleic Acids Res. 2018;46(D1):D1074–D1082. doi:10.1093/nar/gkx1037

31. DrugBank | Powering health insights with structured drug data. Accessed February 6, 2024. https://www.drugbank.com/

32. Drug Group | DrugBank Help Center. Accessed February 6, 2024. https://dev.drugbank.com/guides/terms/drug-group

33. Manning G, Whyte DB, Martinez R, Hunter T, Sudarsanam S. The protein kinase complement of the human genome. Science. 2002;298(5600):1912–1934. doi:10.1126/science.1075762

34. KinBase: Kinase Database at Manning’s Group. Accessed February 6, 2024. http://kinase.com/web/current/kinbase/genes/SpeciesID/9606/

35. Seal RL, Braschi B, Gray K, et al. Genenames.org: the HGNC resources in 2023. Nucleic Acids Res. 2022;51(D1):D1003–D1009. doi:10.1093/nar/gkac888

36. G protein-coupled receptors | HUGO Gene Nomenclature Committee. Accessed February 6, 2024. https://www.genenames.org/data/genegroup/#!/group/139

37. Jassal B, Matthews L, Viteri G, et al. The reactome pathway knowledgebase. Nucleic Acids Res. 2020;48(D1):D498–D503. doi:10.1093/nar/gkz1031

38. Downloads - Reactome Pathway Database. Accessed February 6, 2024. https://reactome.org/download-data

39. Aebersold R, Agar JN, Amster IJ, et al. How many human proteoforms are there? Nat Chem Biol. 2018;14(3):206–214. doi:10.1038/nchembio.2576

40. Peters JU. Polypharmacology – Foe or Friend? J Med Chem. 2013;56(22):8955–8971. doi:10.1021/jm400856t

41. Mullard A. FDA approves first-in-class SYK inhibitor. Nat Rev Drug Discov. 2018;17(6):385–385. doi:10.1038/nrd.2018.96

42. Paik J. Fostamatinib: A Review in Chronic Immune Thrombocytopenia. Drugs. 2021;81(8):935–943. doi:10.1007/s40265-021-01524-y

43. Fabregat A, Sidiropoulos K, Viteri G, et al. Reactome pathway analysis: a high-performance in-memory approach. BMC Bioinformatics. 2017;18(1):142. doi:10.1186/s12859-017-1559-2

44. Ortiz G, Kutateladze TG, Fujimori DG. Chemical tools targeting readers of lysine methylation. Curr Opin Chem Biol. 2023;74:102286. doi:10.1016/j.cbpa.2023.102286

45. Wu Q, Heidenreich D, Zhou S, et al. A chemical toolbox for the study of bromodomains and epigenetic signaling. Nat Commun. 2019;10(1):1915. doi:10.1038/s41467-019-09672-2

46. Scheer S, Ackloo S, Medina TS, et al. A chemical biology toolbox to study protein methyltransferases and epigenetic signaling. Nat Commun. 2019;10(1):19. doi:10.1038/s41467-018-07905-4

47. Wang J, Yazdani S, Han A, Schapira M. Structure-based view of the druggable genome. Drug Discov Today. 2020;25(3):561–567. doi:10.1016/j.drudis.2020.02.006

48. Drewry DH, Wells CI, Andrews DM, et al. Progress towards a public chemogenomic set for protein kinases and a call for contributions. PLoS ONE. 2017;12(8):e0181585. doi:10.1371/journal.pone.0181585

49. Roth BL, Irwin JJ, Shoichet BK. Discovery of new GPCR ligands to illuminate new biology. Nat Chem Biol. 2017;13(11):1143–1151. doi:10.1038/nchembio.2490

50. Knapp S, Arruda P, Blagg J, et al. A public-private partnership to unlock the untargeted kinome. Nat Chem Biol. 2013;9(1):3–6. doi:10.1038/nchembio.1113

51. Lahiry P, Torkamani A, Schork NJ, Hegele RA. Kinase mutations in human disease: interpreting genotype–phenotype relationships. Nat Rev Genet. 2010;11(1):60–74. doi:10.1038/nrg2707

52. Heng BC, Aubel D, Fussenegger M. An overview of the diverse roles of G-protein coupled receptors (GPCRs) in the pathophysiology of various human diseases. Biotechnol Adv. 2013;31(8):1676–1694. doi:10.1016/j.biotechadv.2013.08.017

