## Supplementary Methods & Supplementary Figure 1 for "Chemical Coverage of the Human Reactome"

Supplementary Methods........................p.2

Supplementary Figure 1..........................p.4

### Supplementary Methods

***Data collection: compound libraries, protein families, and human biological pathways***

*Chemical Probes*

In total, 554 chemical probes that target human proteins were compiled from Chemical Probes Portal[14,22], which includes probes developed by the SGC, opnMe[13,23], and Donated Chemical Probes[24,25]. For probes sourced from Chemical Probes Portal, only the ones given an in-cell rating of 3 or more were included.[14]

*Chemogenomics Compounds*

484 chemogenomic compounds, consisting of 410 kinase ligands and 74 GPCR ligands, were compiled from: the kinase chemogenomic set (KCGS) V2.0[26,27] released by the SGC at the University of North Carolina at Chapel Hill and the EUbOPEN chemogenomic set[28], consisting of a protein kinase and GPCR family subset at the time of this study.

*Drugs*

Version 5.1.11 of all approved small molecule drugs that target human proteins (1693) were downloaded from DrugBank.[29–31] DrugBank defines an approved drug as one that has been approved in at least one jurisdiction at some point in time.[32] DrugBank provides extensive data on drug targets, listing all targets that have been found to be involved in the physiological effect, including those that play a role in the transport or activation of the drug.[29]

*Human Kinases and GPCRs*

A list of 538 human protein kinases was sourced from KinBase provided by Manning's group[33,34] and a list of 802 human GPCRs was sourced from the HUGO Gene Nomenclature Committee[35,36].

*Human Biological Pathways*

Human biological pathways and human proteins that belong in each pathway were extracted from the Reactome knowledgebase.[37,38]

***Mapping of compounds onto proteins, protein families, and human Reactome pathways***

Compounds were mapped onto proteins, protein families, and human Reactome pathways using the Uniprot ID of the target proteins. A list of all 2,331 proteins targeted by the compounds included in our analysis, as well as the number of chemical probes, chemogenomic compounds, and/or drugs that target them can be found in Table S1. For each of the human kinase and GPCR families, the number of chemical probes, chemogenomic compounds, and drugs that target them can be found in Table S3 and S4 respectively. For mapping of compounds onto human Reactome pathways, separate mappings were conducted using all 11,175 human proteins (Table S2), the 362 human kinases (Table S5), and the 705 human GPCRs (Table S6) that are found in the Reactome pathways.

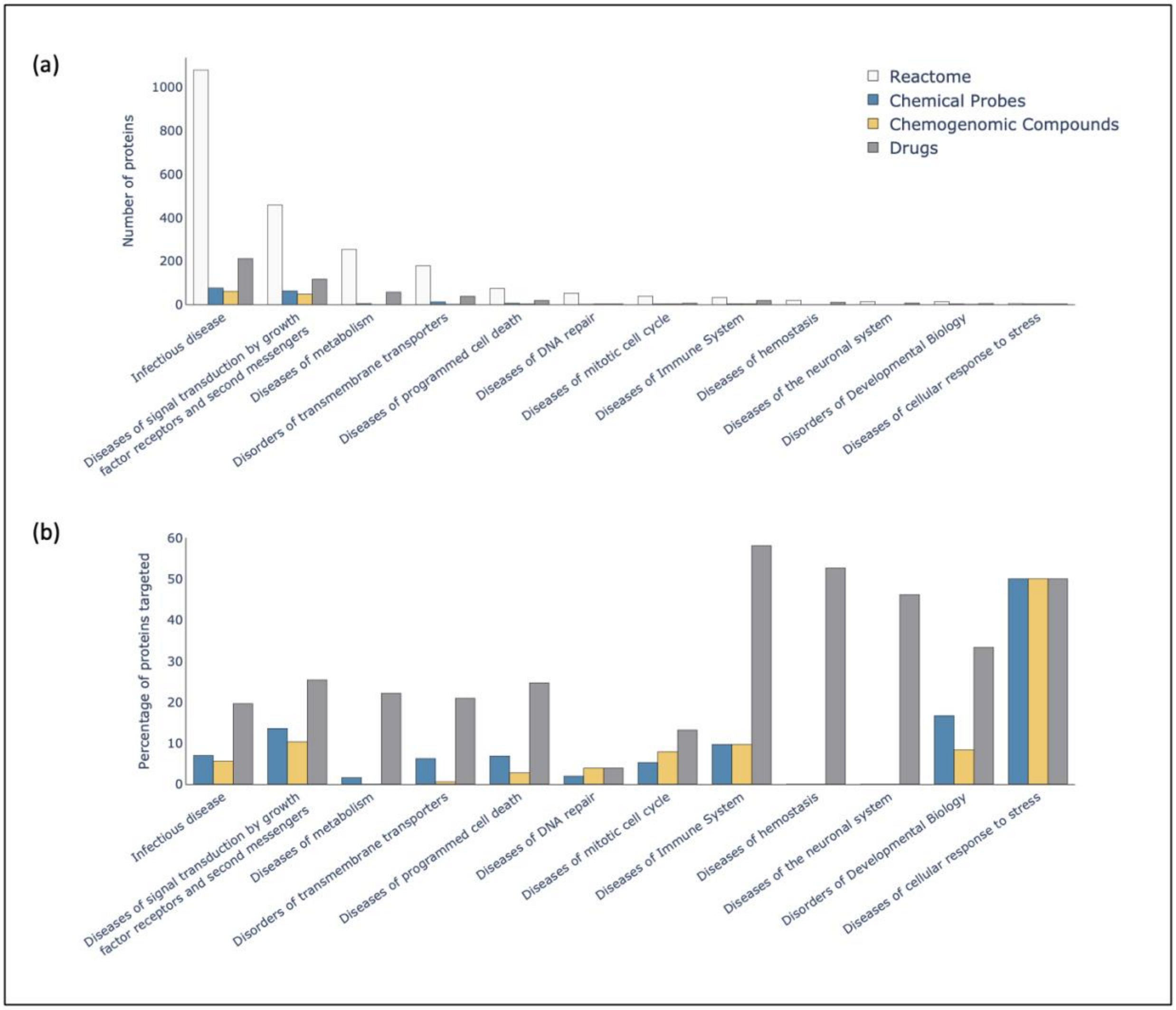


**Figure S1. Chemical coverage of disease pathways.** For each pathway, the number of proteins (a) and % of proteins (b) targeted by chemical probes, chemogenomic compounds and drugs are shown.
